# BRDKRM: An Explainable Framework for Disease Modifying Drug Identification

**DOI:** 10.1101/2024.09.24.614653

**Authors:** Aishik Chanda, Ashmita Dey, Mrittika Chakraborty, Utsav B. Maulik, Sanghamitra Bandyopadhyay

## Abstract

Drug classification into disease-modifying (DM) and symptomatic (SYM) categories is crucial for clinical decision-making and therapeutic strategy development. To address the limitations of current methods, which often lack transparency and interpretability, we propose the Boundary Restricted Dynamic Key Route Mapping (BRDKRM) framework. This novel approach leverages the contextual overlap between disease and drug nodes in a heterogeneous graph, aggregating genes from the top K shortest paths to delineate disease neighborhood boundaries. Inspired by the classic Hansel and Gretel folklore, BRDKRM metaphorically marks boundary nodes along metapaths from disease to drug, akin to Hansel‘s breadcrumbs, which are then used to classify the therapeutic effect of candidate drugs. Our method achieved 86.78% accuracy in categorizing drug-disease treatments and identified 530 genes involved in both disease modification and symptomatic relief. The efficacy of BRDKRM is demonstrated through case studies on multiple sclerosis, offering an explainable approach to drug classification that bypasses extensive clinical trials. By providing biologically sound interpretations of drug classifications, our framework enhances understanding of therapeutic interventions, paving the way for more precise and efficient healthcare solutions while offering a novel approach to mapping disease-drug interactions.

## 1 Introduction

Drug classification plays a crucial role in modern medicine, with the recent focus shifting towards long-term disease modification rather than short-term symptom improvement [9]. Disease-modifying (DM) drugs aim to alter disease progression by targeting root causes or mechanisms, while symptomatic (SYM) drugs provide temporary relief without affecting the underlying disease course [20]. This distinction is vital for clinical trials, patient stratification, and drug safety studies [5, 7].

The categorization of drugs as DM or SYM has significant implications for research and patient care. DM drugs often focus on long-term outcomes and biomarkers of disease progression, while SYM drugs emphasize immediate symptom relief. This classification allows researchers to define relevant endpoints and helps in patient stratification for clinical trials at different disease stages. However, it ‘s important to note that DM drugs may come with more severe adverse effects compared to SYM drugs, necessitating careful consideration and monitoring of their use [28, 32].

The advent of high-throughput technologies and large-scale biological datasets has revolutionized drug discovery and classification. Network-based approaches have emerged as powerful tools for analyzing complex biological systems, effectively prioritizing disease-causing genes and predicting drug-disease associations [1, 24, 30]. These methods represent genes, proteins, drugs, and diseases as nodes in a network, with their interactions as edges, capturing the intricate relationships within biological systems.

Recent computational methods have focused on drug repositioning, expression profile mapping, and machine learning techniques for predicting drug indications [11, 12, 22]. These include data-driven similarity measures, knowledge graphs, and graph convolutional networks. While these approaches have shown promise, they often lack interpretability, a crucial aspect in biomedical research where understanding the reasoning behind predictions is essential.

To address this gap, we present the Boundary Restricted Dynamic Key Route Mapping (BRDKRM) framework. This novel approach exploits contextual overlap between disease and drug nodes within a heterogeneous graph structure to classify drugs as DM or SYM for specific conditions. BRDKRM employs a modified walk algorithm to determine drug-disease associations, providing explainable insights by identifying significant genes associated with drug indication classification.

Our method aggregates genes from top K shortest paths connecting diseases and known drugs, delineating disease neighborhood boundaries. We then classify novel drug-disease interactions based on average scores of boundary genes within paths leading to unidentified drugs. This approach offers a biologically grounded and interpretable drug classification, potentially reducing the need for extensive clinical trials. It enhances the understanding of a drug‘s potential impact on disease progression and aids in identifying common genetic markers to elucidate drug indications effectively.

By providing a more nuanced understanding of drug indications, BRDKRM can guide researchers and clinicians in making informed decisions about therapeutic interventions. This is particularly valuable in the context of complex diseases like neurodegenerative disorders, where the distinction between DM and SYM drugs can significantly impact treatment strategies [6, 8].

## 2 Materials and Method

### 2.1 Construction of Heterogeneous Biological Graph

We constructed a heterogeneous graph to enhance our study, integrating diverse biological datasets from public resources. The graph‘s metanodes represent complex diseases, drugs, disease-associated genes, and drug targets. Diseases were selected for clinical relevance, ensuring specificity while allowing comprehensive annotation from the Disease Ontology (DO) database [39]. Protein-coding human genes implicated in these diseases and drug targets were curated from Entrez Gene [23]. Approved small molecule compounds were obtained from DrugBank [19]. Edges capture relationships between node entities. To ensure biologically meaningful disease-drug paths, we restricted paths to include only genes, pathways [36–38], biological processes, cellular components, and molecular functions [40]. We reversed outgoing edges from drug nodes and converted all directed edges to undirected, preventing drugs from acting as intermediaries in paths. Relationships between drug classifications and diseases were curated from PharmacotherapyDB [13]. This comprehensive graph integrates multiple biological data types, facilitating the exploration of complex relationships between diseases, drugs, and their molecular targets, while preserving the integrity of our path-based analytical approach.

### 2.2 BRDKRM: Proposed Framework

The premise of our study is based on the assumption that the overlap in the context of a disease node and a drug node can determine whether a drug behaves as a disease-modifying drug or a symptomatic drug for a specific disease. Leveraging this assumption, we propose a novel framework that extracts the overlapping context between a disease and a drug to classify the drug as either disease-modifying or symptomatic based on this context. The pipeline of the proposed method is illustrated in Figure 1a, while the detailed graphical representation of the algorithm is depicted in Figure 1b.

**Fig 1.**
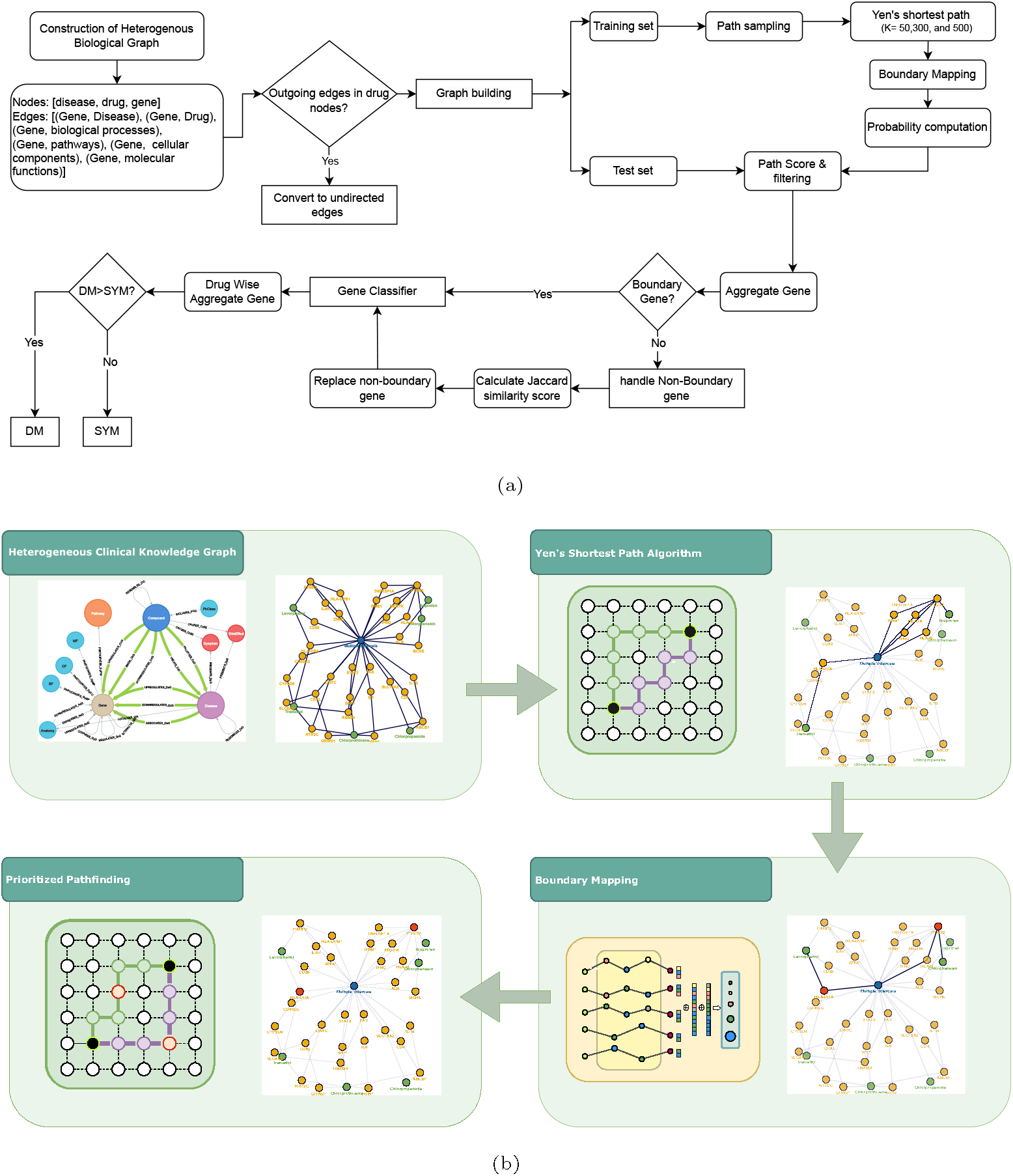
The figure illustrates (a) the complete pipeline and (b) an abstract overview of our framework BRDKRM.

#### 2.2.1 Boundary Mapping

Our dataset comprised disease, drug, and their relationship types DM or SYM. For processing the training set as explained in Algorithm 1, we began by listing all drugs associated with each disease and computing the *k*-shortest paths between each disease and its corresponding drugs using Yen‘s shortest path algorithm [35]. We sampled the paths from the graphs into *n* paths where *n* = 50, 300, and 500 paths between a disease and a drug. We aggregated the genes in these paths and calculated the frequency of each gene‘s occurrence in paths associated with disease-modifying and symptomatic drugs. Each gene was thus assigned two frequencies: one for disease-modifying and one for symptomatic.

For the test set, we again listed all drugs for each disease and computed the *k*-shortest paths between them as explained in Algorithm 2. We then compiled a list of known genes for each disease based on the training set. For each gene, we calculated the probability of it being associated with either category using the formula:

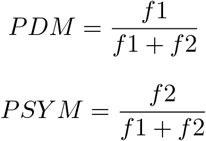

where *f* 1 and *f* 2 are the frequencies of the gene in disease-modifying and symptomatic drug paths, respectively. The confidence for each gene was determined as the maximum of PDM and PSYM. We then sorted the paths for each disease-drug pair, prioritizing paths with the most known genes and those where the confidence of the genes exceeds a threshold *T*. From this sorted list, we selected the top *N* paths and aggregated all the genes within them, marking genes not present in the known set as unknown.

##### Algorithm 1

Training Set Processing

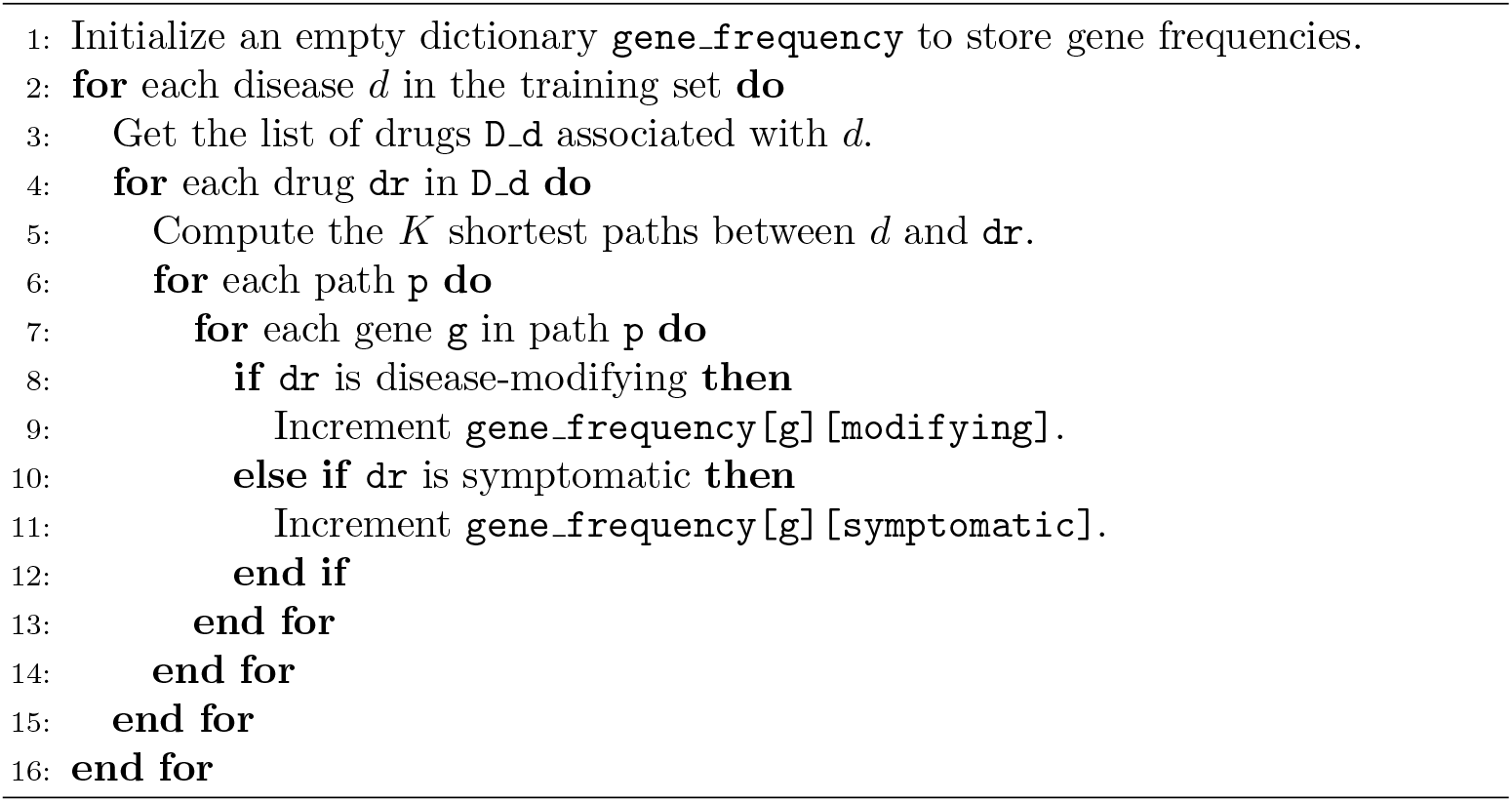

##### Algorithm 2

Test Set Processing

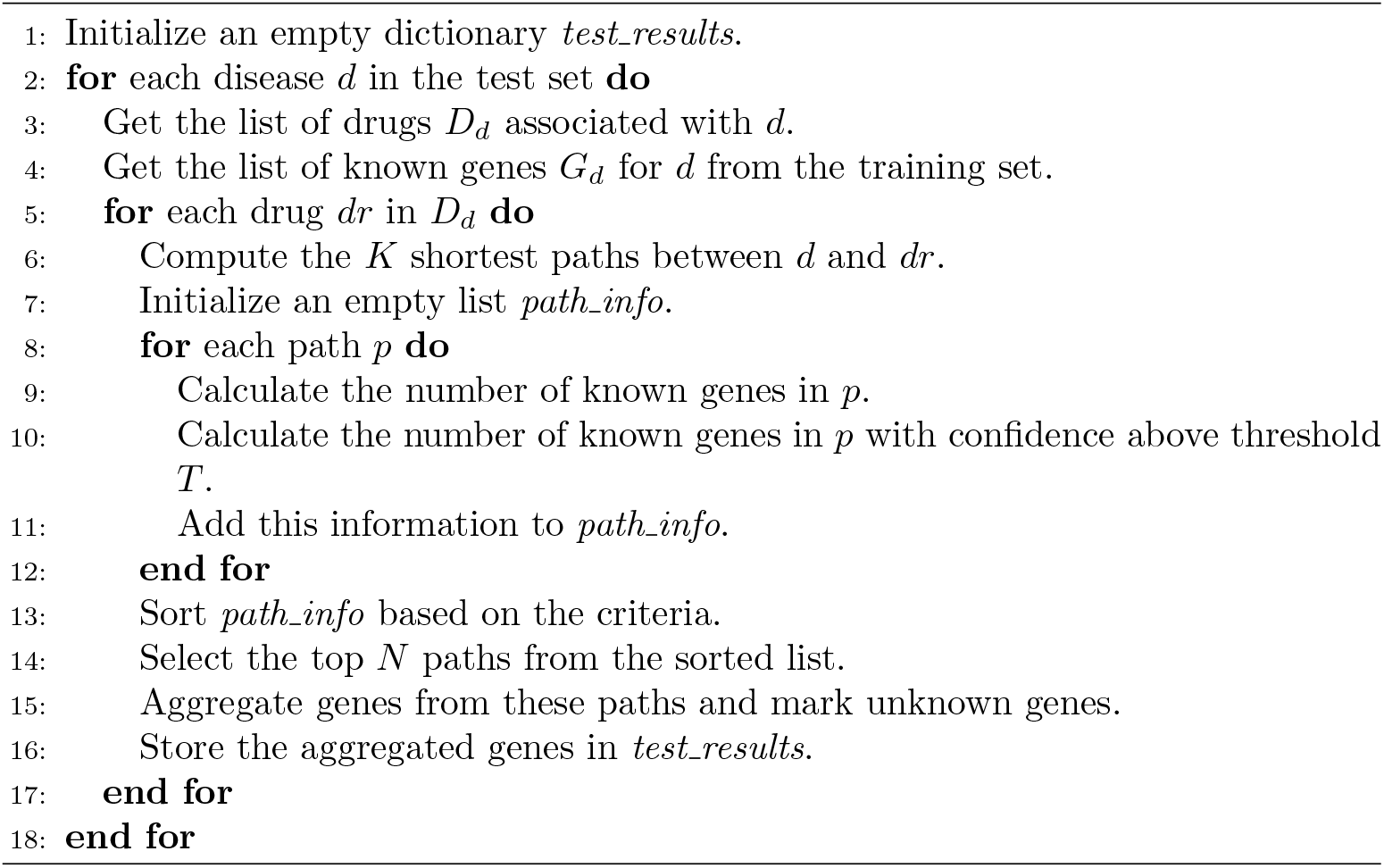

To address the issue of unknown genes, we computed node similarity between all possible pairs in the graph. This strategy is explained in Algorithm 3. The similarity has been calculated for two nodes based on their neighborhood sets using Jaccard similarity index [3]. To calculate the Jaccard similarity between two nodes *u* and *v* in a graph, we use the following formula:

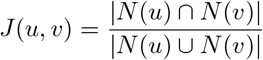

where:*N* (*u*) is the set of neighbors of node *u* and *N* (*v*) is the set of neighbors of node *v*. |*N* (*u*) ∩ *N* (*v*) is the number of common neighbors between nodes *u* and *v* while *N* (*u*) ∪ *N* (*v*) is the number of unique neighbors of nodes *u* and *v*.

For each unknown gene, we identified the top 10 most similar genes and checked if any of these were known genes. For a high similarity score, if any node satisfied this condition, we replaced the unknown gene with the known gene. This step aims to enhance the accuracy of our gene-based context determination by leveraging similarity-based inference.

##### Algorithm 3

Fixing the Unknown Genes

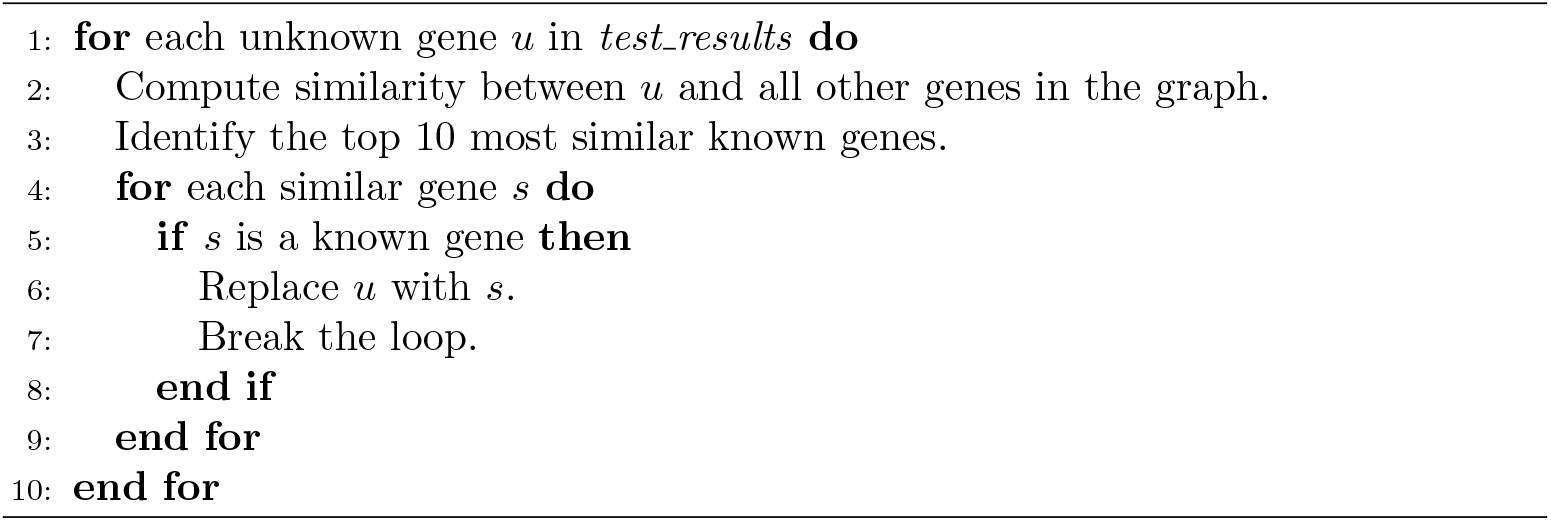

#### 2.2.2 Classifying Drug Categories

We classified each disease-drug pair by considering the aggregate list of genes from the top *N* paths. We counted the number of genes with non-zero frequencies for DM and SYM categories. The classification of the disease-drug relationship was based on the category with the greater count of genes explained in Algorithm 4. This comprehensive approach allowed us to leverage the contextual overlap between disease and drug nodes to accurately classify drugs, providing a robust framework for drug repurposing and therapeutic strategy development.

##### Algorithm 4

Classifying Drug Categories

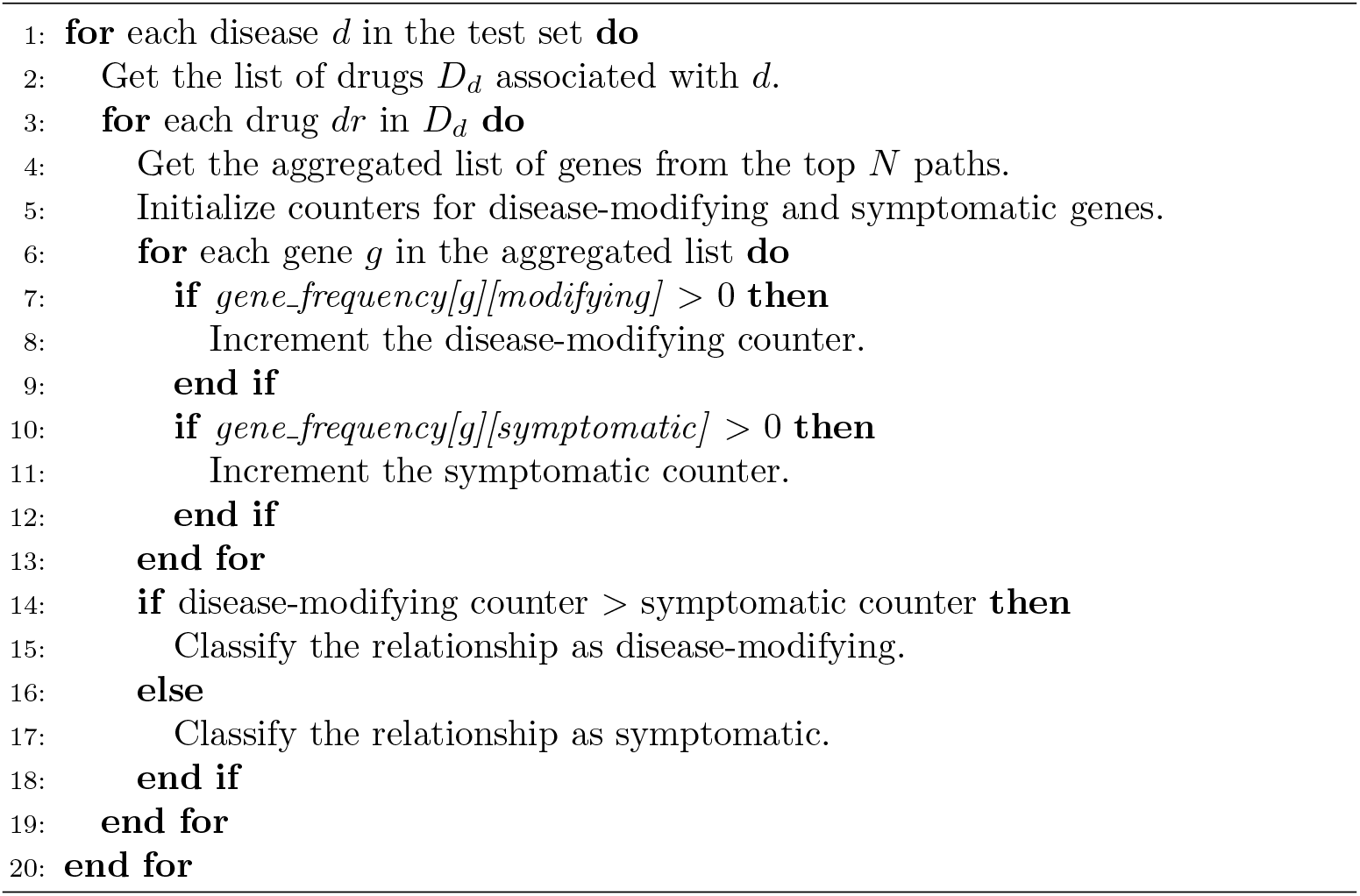

## 3 Results and Discussion

Our study aims to classify drug categories based on context-based overlapping genes. In the growing era of healthcare, advanced machine learning techniques have often failed to decipher the biological significance of molecular entities. In our proposed method, we utilized advanced statistical methods to understand the contribution of genes and biological pathways in the underlying mechanism of drug-disease relationships. Moreover, these methods help predict the treatment procedures by categorizing drugs in DM (disease-modifying) and SYM (symptomatic) categories.

In the proposed framework, Yen‘s shortest path algorithm is applied to compute the *k*-shortest paths for each disease-drug pair. This step is crucial in mapping out the potential routes through which drugs interact with disease nodes. The computation resulted in a comprehensive set of paths that provided a foundation for subsequent analyses. Subsequently, to address the genetic aspects of disease-drug interactions, we computed the node similarity for all possible pairs in the graph. This analysis considered the genetic context of diseases by evaluating the aggregate list of genes from the selected paths. This genetic context provided deeper insights into the mechanisms of action of drugs and their potential effects on different diseases. The similarity scores helped distinguish between disease-modifying drugs and symptomatic treatments, enhancing the accuracy of our classification. A network in Figure 2 shows the associations among diseases based on calculated similarity scores. In this network, nodes represent diseases, and the edges depict their similarities. Furthermore, we identified the overlapping genes between DM and SYM-specific genes along with the pattern genes. These 530 genes play roles in both modifying the disease and providing symptomatic relief, depending on the context of their involvement in specific pathways. The common genes are shown through a Venn diagram in Figure 3A. This high level of precision underscores the reliability of our approach and its potential for widespread application in drug repurposing and therapeutic strategy development. In Figure 3B, shankey plot is performed to validate our a classified drug categories through literature resources.

**Fig 2.**
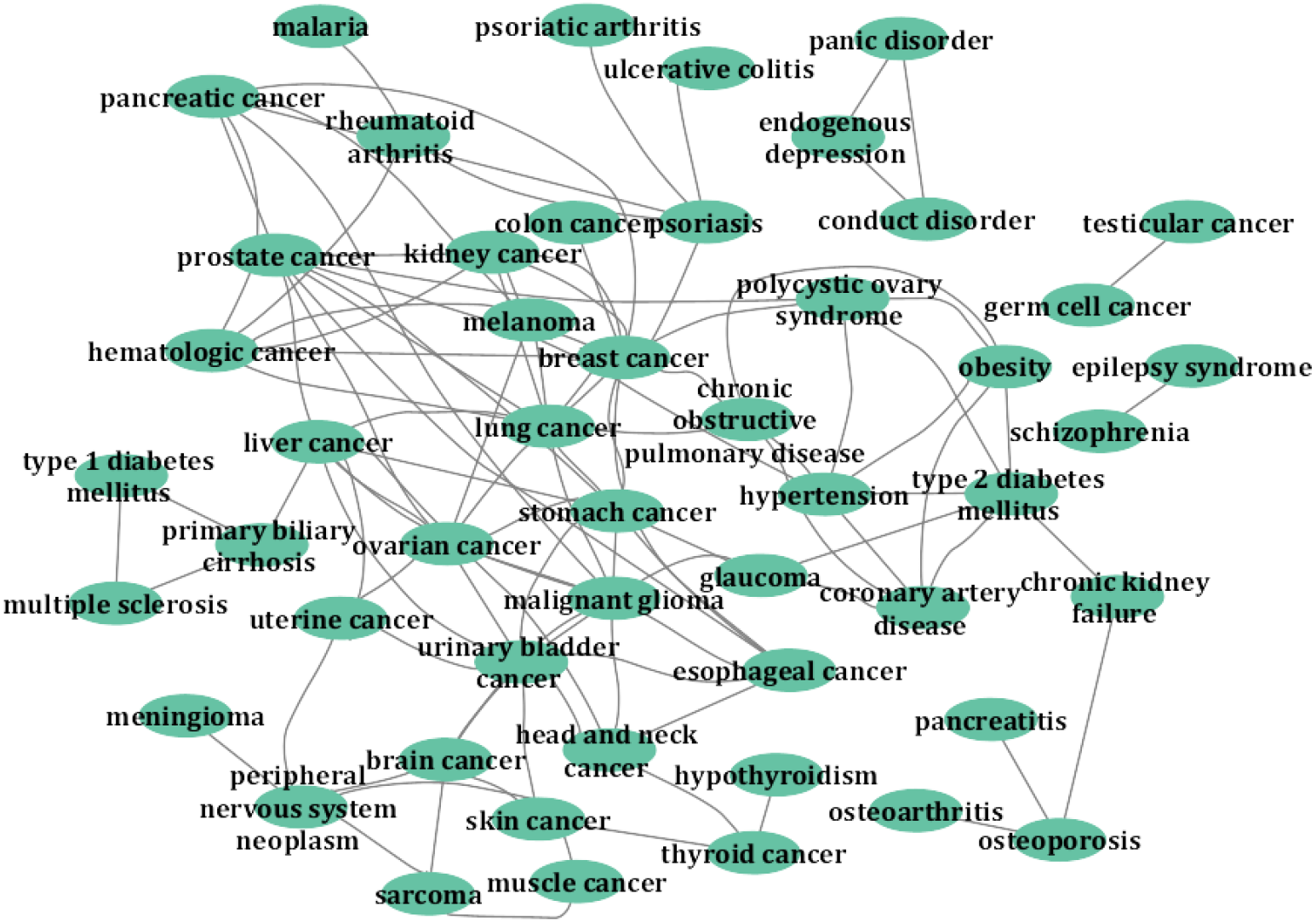
The network analysis we conducted, which maps diseases based on the similarity of their neighborhoods, provides compelling empirical evidence for a novel approach to understanding drug-disease interactions. By visualizing these relationships, we demonstrate that a disease‘s neighborhood can serve as a distinctive characteristic. This finding lends support to our hypothesis that the extent of overlap between a disease node‘s context and a drug node‘s context can be a determinant factor in predicting whether a drug will act as a disease-modifying agent or a symptomatic treatment for a specific condition. This insight opens new avenues for drug discovery and repurposing strategies, potentially revolutionizing our approach to therapeutic interventions across a wide spectrum of diseases.

**Fig 3.**
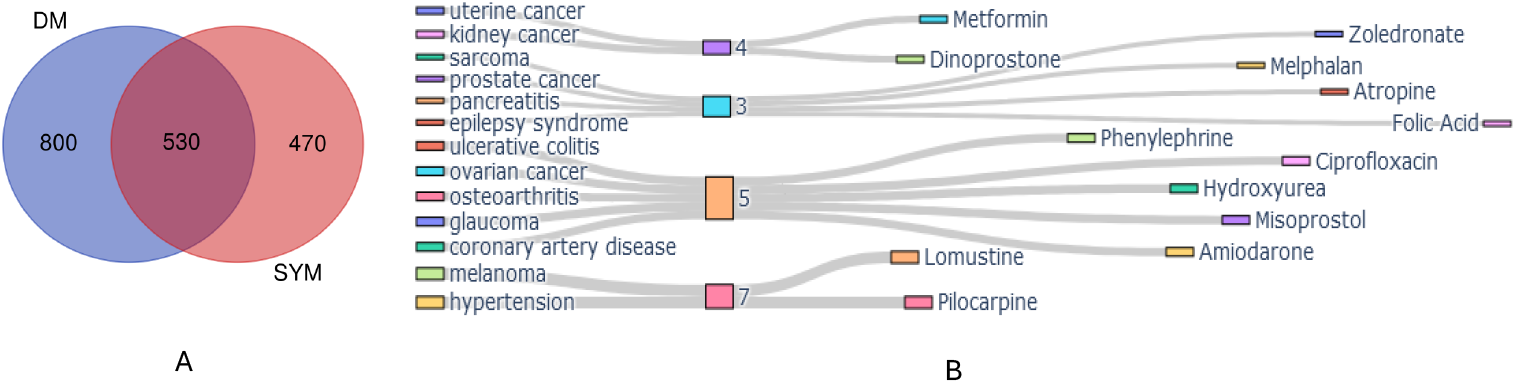
The desired results of our proposed frameworks are shown in (A)the Venn diagram represents the pattern genes responsible for DM and SYM genes and 530 common genes responsible for both the nature of the drugs and (B) the Shankey plot supports our predicting results based on literature review.

### 3.1 Practical Implications

While the study focuses on drug categorization, our network analysis also revealed significant opportunities for drug repurposing. By identifying drugs with strong associations with multiple diseases, our framework highlights potential candidates for repurposing. As an example, we have found a significant amount of both categories of DM and SYM drugs in the case of multiple sclerosis. However, detailed studies are noted in the literature [25], making this disease ideal to demonstrate and validate our proposed framework. In this regard, a network has been established that is dedicated to this disease along with its drug categories and targeted genes. Our method distinctly categorizes the drugs into two categories and deciphers the pattern genes responsible for the categorization. The network is shown in Figure 4. The red node depicts the disease itself, while genes, DM, and SYM drugs are represented through brown, purple, and yellow nodes respectively. The accurate classification of drugs for multiple sclerosis aids in clinical decision-making by providing a clearer understanding of which drugs modify the disease course and which merely alleviate symptoms. This distinction is vital for developing effective treatment plans, and ensuring that patients receive therapies that offer the most benefit, especially for diseases where medication options are limited due to a lack of knowledge regarding the disease mechanism.

**Fig 4.**
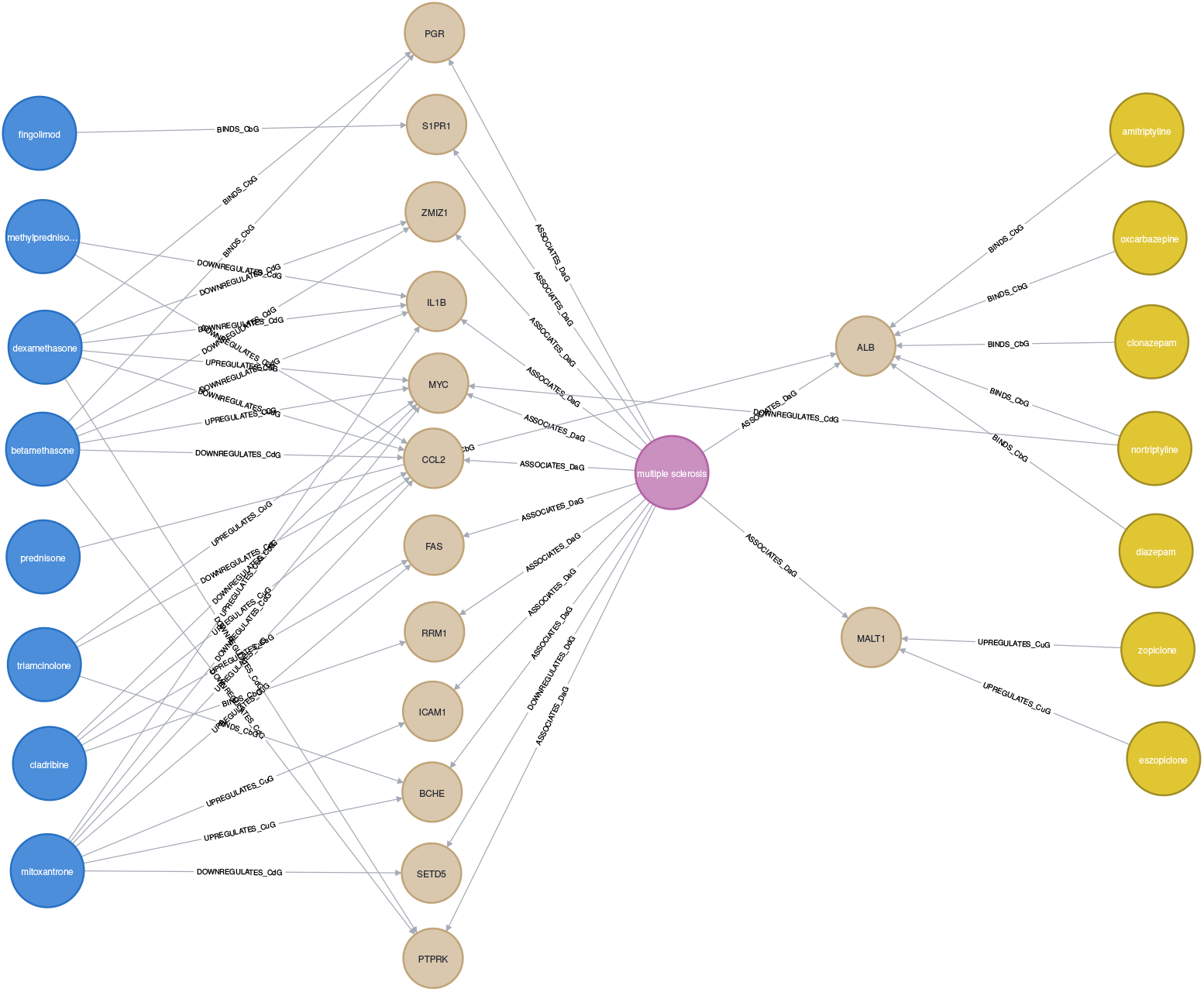
BRDKRM effectively classifies drugs into two distinct categories and identifies the gene patterns responsible for this classification. In the network, we selected multiple sclerosis as a disease node, represented by the red node, while genes, DM drugs, and SYM drugs are depicted by yellow, blue and purple nodes, respectively.

**Fig 5.**
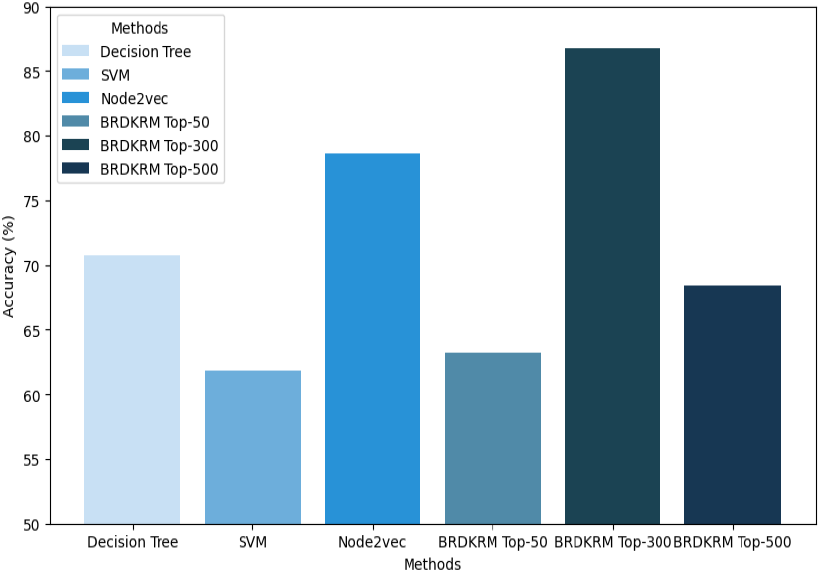
Comparison of Model Accuracy Across Different Algorithms: The bar graph illustrates the performance of various machine learning models, including Decision Tree [41], SVM [42], Node2vec [43], and three configurations of the BRDKRM model. The BRDKRM Top-300 configuration achieves the highest accuracy, highlighting the importance of selecting the right parameters for optimal performance.

### 3.2 Comparative Analysis

To rigorously assess the performance of our proposed method, we conducted a series of experiments comparing it against several established machine learning models, including Random Forest (RF), Support Vector Machine (SVM), and Graph Neural Network (GNN). These models are chosen for their relevance and widespread use in similar tasks. We applied our method, alongside the RF, SVM, and GNN models, to the same dataset to ensure a fair comparison.

**Decision Tree [41]:** Decision Trees are a popular choice for classification tasks due to their simplicity and interpretability. However, they often struggle with high-dimensional data and may suffer from overfitting.

**Support Vector Machine [42]:** SVM performed well in cases where the decision boundary was clear, but it was less effective in handling non-linear relationships without extensive kernel tuning, which impacted its overall accuracy.

**node2vec [43]**: node2vec is a graph-based embedding technique that learns low-dimensional representations for nodes in a graph. This method is particularly useful in capturing the structural properties of the data.

Each model is fine-tuned to its optimal hyperparameters using cross-validation, and the accuracy score is used as the primary evaluation metric. The experiments are conducted in a controlled environment, with consistent preprocessing, feature selection, and training procedures.

The comparison revealed that our proposed method outperformed the Decision Tree, SVM, and GNN models in terms of accuracy. Specifically, our method achieved an accuracy score of 86.78%, compared to 70.79% for the Decision Tree model, 61.80% for the SVM model, and 78.65% for the node2vec model.

## 4 Conclusion

BRDKRM introduces a transformative approach in healthcare by offering an interpretable method to gather information on repurposed drugs, bypassing the extensive clinical trial process. With the surge in methods and data, domain experts often struggle to decode the biological significance of advanced learning techniques. Due to a lack of domain explainability most of the designed pipelines failed to provide the desired output. Our framework addresses this by accurately classifying drugs listed as no-indication in public databases and providing biologically sound interpretations of these classifications. The proposed framework is a key solution to increase the efficacy of clinical trial outcomes and enhance the understanding of therapeutic interventions, paving the way for more precise and efficient healthcare solutions.

## Acknowledgements

The research work of MC and SB is supported by the Indo-French Centre for the Promotion of Advanced Research, Ministry of Science and Technology, Govt of India through the project: 6702-1‘Exploring Graph Neural Networks (GNN‘s) for DATA-Driven Modeling of Poly-Pharmacy‘. In addition, SB acknowledges the JC Bose Fellowship grant No. JBR/2021/000036 from SERB, Govt of India.

